# PromptBio-Bench: Benchmarking LLM-based Bioinformatics Agents for End-to-End Data Analysis

**DOI:** 10.64898/2026.05.05.723092

**Authors:** Wenbin Guo, Minzhe Zhang, Bowei Han, Youjia Ma, Yang Leng, Shishir Hebbar, Xiaoyuan Zhou, Wenhao Gu, Xiao Yang, Shashi Dhar

## Abstract

Large language model (LLM)-based agents hold transformative potential for automating bioinformatics workflows; however, systematic evaluations of their capabilities remain limited, hindering a clear assessment of their readiness for real-world application. We introduce PromptBio-Bench, a comprehensive evaluation suite of 244 expert-curated tasks spanning bioinformatics and data science at varied difficulty levels, and an evaluation framework for structured file comparison and scoring against expert reference answer files. Evaluation of three state-of-the-art bioinformatics agents revealed comparable performance between Biomni and ToolsGenie, with all agents showing a marked decline in accuracy as task difficulty increased. As foundation models and agent frameworks continue to evolve, PromptBio-Bench provides a valuable benchmark infrastructure for systematically tracking progress in agentic bioinformatics.

## Introduction

The exponential growth of biomedical data^1^, driven by high-throughput omics technologies, has created an urgent need for scalable, reproducible bioinformatics analysis. Traditional approaches rely heavily on human expertise in computational environment management, manual tool configuration, and specialized programming skills, imposing significant barriers for researchers without computational training while also constraining analytical throughput even for experienced bioinformaticians. The emergence of agentic bioinformatics^2^, powered by large language model (LLM)-based agents, offers a transformative path toward more accessible, scalable, and reproducible bioinformatics analysis. By integrating natural language reasoning with autonomous code generation, dynamic tool use, and iterative self-refinement, these agents have the potential to lower technical barriers and accelerate the conversion of raw biomedical data into biological insights^3^.

Recent years have witnessed rapid progress in AI agents for bioinformatics, ranging from domain-specialized systems to workflow-oriented agents and increasingly general-purpose assistants^4^. Domain-scoped agents have shown promise in focused biological settings. For example, CellVoyager^5^ and SpatialAgent^6^ automate single-cell and spatial transcriptomics analysis, respectively; DrBioRight 2.0^7^ enables large-scale exploration of cancer proteomics; CRISPR-GPT^8^ focuses on gene-editing experiment design; and PathChat^9^ and SPARK^10^ support pathology-centered multimodal reasoning and discovery. Beyond these specialized applications, Early workflow-oriented systems such as AutoBA^11^ and BioMaster^12^ demonstrated the feasibility of LLM-driven automation for multi-omic bioinformatics analysis, with AutoBA focusing on code generation and execution for omics workflows and BioMaster extending this direction through retrieval-guided planning, debugging, and output validation. More recent systems further broaden the agentic paradigm by expanding how agents access, organize, and adapt their action spaces. Biomni^13^ integrates a broad, curated biomedical action space spanning domain tools and databases, coupled with ReAct-style planning and execution; STELLA^14^ emphasizes self-evolution, autonomously expanding its reasoning strategies and tool repertoire with experience; ToolsGenie^15^ prioritizes operational robustness through dynamic Docker environment management and modular multi-agent extensibility; and BioMedAgent^16^ focuses on learning, chaining, and reusing bioinformatics tools through interactive exploration and memory retrieval. Collectively, these systems represent encouraging progress toward adaptive, autonomous, and agent-driven bioinformatics analysis.

Despite rapid progress in bioinformatics agents, their evaluation in practical analytical workflows remains limited. Existing benchmarks capture important but partial dimensions of agent capability. LAB-Bench^17^ primarily evaluates biological knowledge and research reasoning, while BixBench^18^ moves closer to practical bioinformatics by introducing expert-curated analytical questions involving biological datasets. BioMysteryBench^19^ and CompBioBench^20^ further advance objective evaluation by emphasizing open-ended computational-biology problem solving with objectively verifiable answers, while complementary efforts such as BiomniBench^21^ evaluate agent trajectories and process quality and BioML-bench^22^ focuses on end-to-end biomedical machine-learning tasks. Together, these benchmarks provide valuable foundations for evaluating biological reasoning, data analysis, agent behavior, and machine-learning workflows. However, they do not fully capture the breadth of real-world bioinformatics practice, where agents must interpret diverse task-specific instructions, operate across heterogeneous biomedical and data-analytic contexts, execute computational workflows, and produce outputs ranging from structured bioinformatics files and tabular results to visualizations and text summaries. This gap motivates the development of a more comprehensive benchmark for day-to-day bioinformatics and data-science analysis.

To address this need, we introduce PromptBio-Bench, a comprehensive benchmarking suite for LLM-based bioinformatics agents. PromptBio-Bench includes 244 tasks designed to reflect the breadth and analytical diversity of bioinformatics and data science, spanning genomics, epigenomics, transcriptomics, proteomics, single-cell omics, data wrangling, statistical modeling, and machine learning. Each task provides a natural language description, input data files, and a reference answer produced by an experienced human bioinformatician, serving as the gold-standard baseline for evaluation. A distinctive feature of PromptBio-Bench is its evaluation framework: rather than relying on multiple-choice answers or simple task-completion flags, it combines pre-built comparison tools for structured bioinformatics file formats, automated quantitative metrics for tabular outputs, and LLM-as-judge scoring for unstructured outputs such as text summaries and images. This design enables fine-grained assessment across heterogeneous task types and output formats.

Using PromptBio-Bench, we evaluated three representative general-purpose bioinformatics agents, Biomni, STELLA, and ToolsGenie, across end-to-end task completion, analytical accuracy, computational runtime, and token consumption. Biomni and ToolsGenie showed broadly comparable performance across the benchmark, completing nearly all tasks with completion rates of 0.99 and 0.98, respectively, whereas STELLA completed 0.88 of tasks. Despite the strong execution robustness of Biomni and ToolsGenie, analytical accuracy remained moderate and declined with increasing task difficulty across all agents, highlighting a persistent gap between successful workflow execution and expert-level analytical correctness. Computational cost further revealed distinct efficiency profiles, with Biomni showing the shortest runtime, ToolsGenie incurring moderate additional overhead, and STELLA requiring substantially greater runtime and token usage. Together, these findings highlight meaningful differences in the trade-offs between execution reliability, analytical performance, and computational efficiency across current bioinformatics agents. By providing a standardized, difficulty-stratified task suite and a rigorous evaluation framework, PromptBio-Bench serves as a valuable community resource for tracking AI agent capabilities in computational biology and guiding the development of more capable and efficient systems.

## Results

### PromptBio-Bench: Benchmark Design and Task Suite

To assess the ability of LLM-based agents to perform real-world computational biology tasks, we developed PromptBio-Bench, a benchmark suite built around standardized task capsules and an structured evaluation framework (Figure 1A). Each task capsule contains a natural-language task description, associated input data files, and a human expert-generated reference answer accompanied by an evaluation guide. During execution, each agent receives the same task prompt and input data, performs the requested analysis, and returns output files together with an execution log. The resulting outputs are evaluated through a standardized pipeline that verifies output validity, determines output format, applies the appropriate comparison strategy, quantifies similarity to the reference answer, assesses answer equivalence, as well as records computational cost. Structured outputs are compared with human expert reference answers using task-specific similarity metrics, whereas text and image outputs are assessed using LLM-as-judge approach followed by human expert review, as detailed in the Methods.

**Figure 1.**
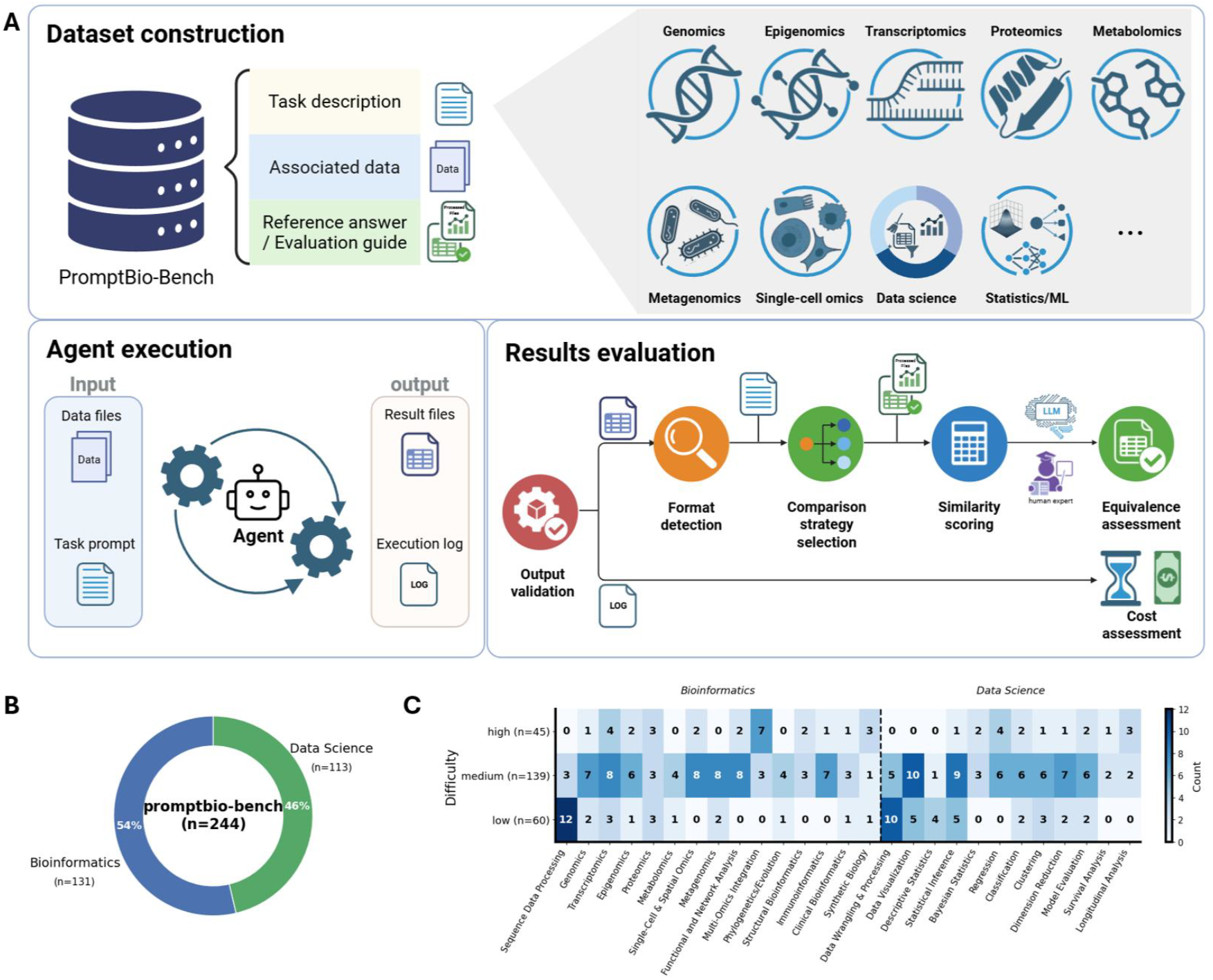
Overview of PromptBio-Bench. (A) *Dataset components (top):* Each benchmark task is organized as a standardized task capsule containing a natural-language task description, associated input data, and an expert-generated reference answer with an evaluation guide. The benchmark spans diverse bioinformatics and data science domains, including genomics, epigenomics, transcriptomics, proteomics, metabolomics, metagenomics, single-cell omics, statistics and machine learning, and other computational analysis areas. *Agent execution pipeline (bottom left):* During agent execution, each agent receives the task prompt and input data files, performs the requested analysis, and returns result files together with an execution log. *Results evaluation pipeline (bottom right):* Agent outputs are then processed through an automated evaluation pipeline that validates output submission, detects output format, selects the appropriate comparison strategy, computes similarity to the reference answer, and assigns an equivalence assessment. Computational cost, including runtime and token consumption, is assessed in parallel. (B) Composition of PromptBio-Bench tasks, including 244 tasks in total, with 131 bioinformatics tasks and 113 data science tasks. (C) Distribution of tasks across subcategories and difficulty levels within the bioinformatics and data science domains, including low difficulty tasks (n = 60), medium difficulty tasks (n = 139), and high difficulty tasks (n = 45).

In total, PromptBio-Bench comprises 244 task capsules spanning two primary domains: bioinformatics (n = 131, 54%) and data science (n = 113, 46%) (Figure 1B). The benchmark captures a broad spectrum of task categories across modern biological research and quantitative analysis, including genomics, epigenomics, transcriptomics, proteomics, metabolomics, metagenomics, single-cell omics, data science, and statistics/machine learning (Figure 1C). Tasks were further stratified into low (n = 60), medium (n = 139), and high (n = 45) difficulty tiers, enabling evaluation of agent performance across varied analytical complexity. Using PromptBio-Bench, we evaluated three representative bioinformatics agents, Biomni, STELLA, and ToolsGenie, under a single-pass protocol, with one attempt per agent per task, to assess their ability to operate across the full breadth of benchmark tasks. For each task and agent, we recorded completion status, equivalence score, computational time, and token consumption, enabling direct comparison of accuracy, robustness, and cost-efficiency across task categories and difficulty levels.

### End-to-End Task Completion Rate

We first assessed each agent’s ability to complete realistic analytical workflows using the end-to-end task completion rate, defined as the proportion of tasks for which an agent produced a valid, non-empty output file within the allotted time. This metric reflects execution robustness across the full task lifecycle, from interpreting natural-language instructions to invoking the required computational steps and producing an evaluable result. It is distinct from analytical accuracy, which was assessed after successful execution. The results showed that the overall completion rates were high across agents, although differences were evident (Figure 2A). Biomni and ToolsGenie achieved near-ceiling completion rates of 0.99 and 0.98, respectively, whereas STELLA reached 0.88. These results indicate that Biomni and ToolsGenie more consistently completed end-to-end workflows across the benchmark. The lower completion rate observed for STELLA suggests that, despite its flexible self-evolving framework, maintaining reliable execution remains a challenge for dynamic agent architectures in heterogeneous analytical settings.

**Figure 2.**
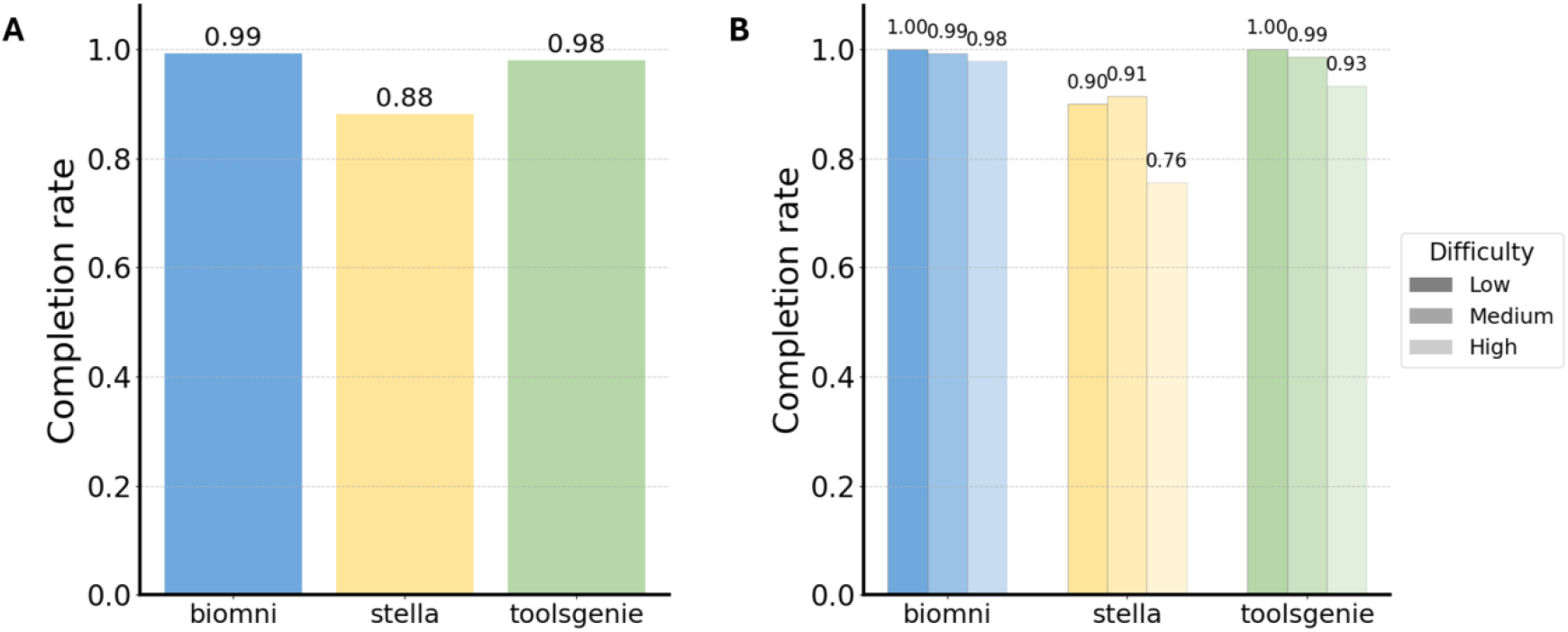
End-to-end task completion rates across agents and difficulty levels. (A) Overall task completion rate for each agent across PromptBio-Bench tasks. (B) Task completion rate stratified by difficulty level (low, medium, high) for each agent.

We next examined whether completion robustness was preserved as task difficulty increased. Biomni maintained consistently high completion rates across low-, medium-, and high-difficulty tasks, with rates of 1.00, 0.99, and 0.98, respectively (Figure 2B). ToolsGenie showed a similar pattern, with completion rates of 1.00, 0.99, and 0.93; the reduction in the high-difficulty tier corresponded to only two incomplete workflows (Supplementary Figure 1). Thus, the high overall completion rates of Biomni and ToolsGenie were largely maintained under increasing analytical complexity. In contrast, STELLA showed greater variability, with completion rates of 0.90, 0.91, and 0.76 across the same difficulty tiers. This pattern suggests that incomplete executions for STELLA were more pronounced among analytically demanding tasks, whereas Biomni and ToolsGenie maintained relatively stable execution across task difficulty levels.

### Output Accuracy Across Tasks

Beyond task completion, we also evaluated the output accuracy by computing similarity scores between each agent-generated output and the corresponding human expert reference answer, with tasks that had failed execution assigned a score of 0. This metric captures the analytical consistency between agent-generated outputs and expert reference answers, jointly reflecting execution success and output quality. Across all tasks, mean accuracy scores were moderate for all three agents (Figure 3A). Biomni and ToolsGenie achieved the same overall accuracy score of 0.76, whereas STELLA scored slightly lower at 0.72. Task-level comparisons further showed that Biomni and ToolsGenie had more concordant correctness profiles with each other than either did with STELLA (Supplementary Figure 2), potentially reflecting differences in their agent architectures and execution strategies. These results indicate that high task completion does not necessarily translate into expert-level analytical accuracy, highlighting task completion and analytical correctness as related but distinct challenges for current bioinformatics agents.

**Figure 3.**
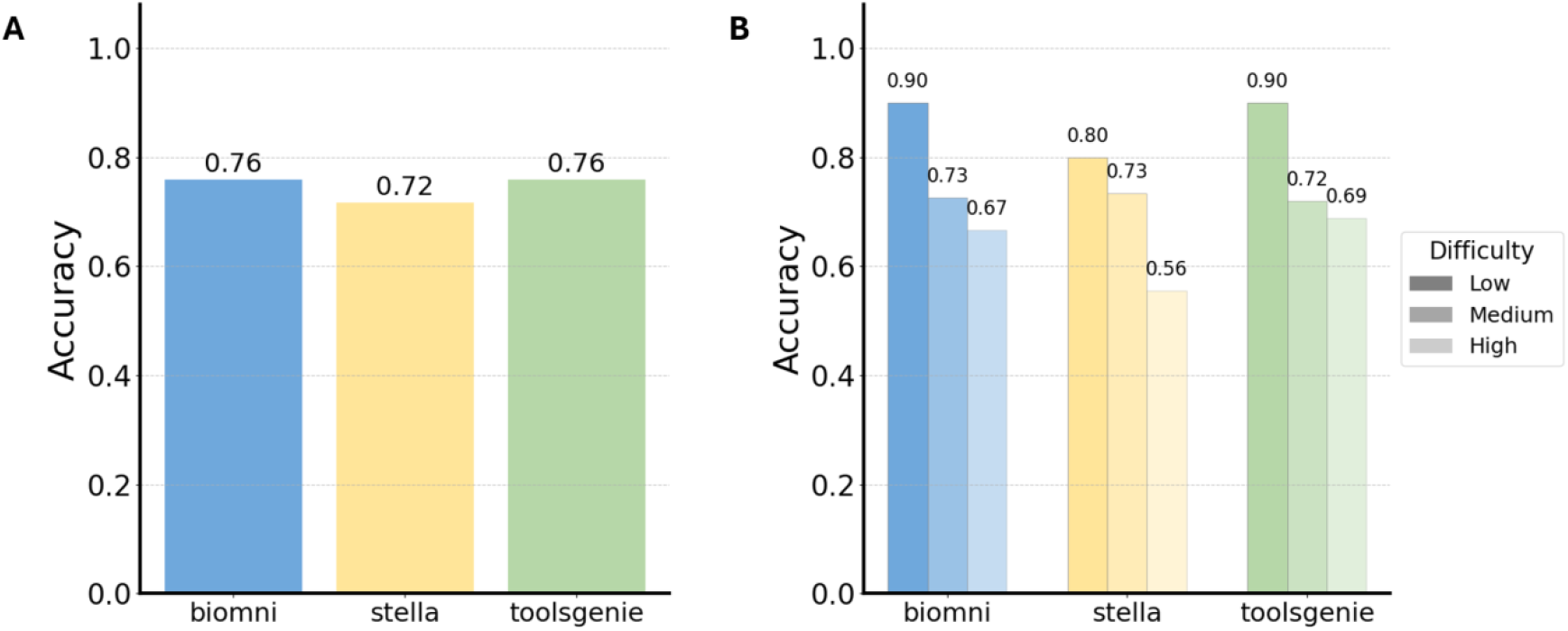
Accuracy across agents and difficulty levels. (A) Mean accuracy score for each agent across all tasks, measured as similarity to human expert reference answers. Failed tasks were assigned a score of 0 and included in the calculation. (B) Mean accuracy scores stratified by task difficulty level.

Accuracy declined with increasing task difficulty across all agents (Figure 3B). On low-difficulty tasks, Biomni and ToolsGenie achieved the highest accuracy scores, both reaching 0.90, while STELLA achieved 0.80. At medium difficulty, the three agents performed similarly, with accuracy scores of 0.73, 0.73, and 0.72 for Biomni, STELLA, and ToolsGenie, respectively. The largest divergence emerged among high-difficulty tasks: ToolsGenie maintained the highest accuracy at 0.69, followed by Biomni at 0.67, whereas STELLA declined to 0.56. This pattern suggests that increasingly complex workflows reduced analytical accuracy across all systems, with the strongest effect observed for STELLA. Together, these findings show that accurately reproducing expert-level outputs on complex bioinformatics workflows remains a central challenge for current agentic systems, even when end-to-end task completion rates are high.

### Runtime and Token Consumption

We next compared the computational cost of each agent in terms of wall-clock runtime and token consumption. Runtime distributions differed across agents (Figure 4A, Supplementary Table 1a). Biomni showed the shortest average runtime and a compact interquartile range, indicating consistent execution across tasks. ToolsGenie showed a moderately higher average runtime and a broader distribution, suggesting that a subset of tasks required longer execution. This pattern may reflect additional overhead from its multi-agent architecture, including agent coordination, result verification, and retry mechanisms. STELLA also showed broad runtime variability, consistent with the additional computational steps required for iterative refinement and multi-model orchestration. These differences indicate that agent architecture can materially affect execution time and should be carefully considered when designing systems for scalable bioinformatics automation.

**Figure 4.**
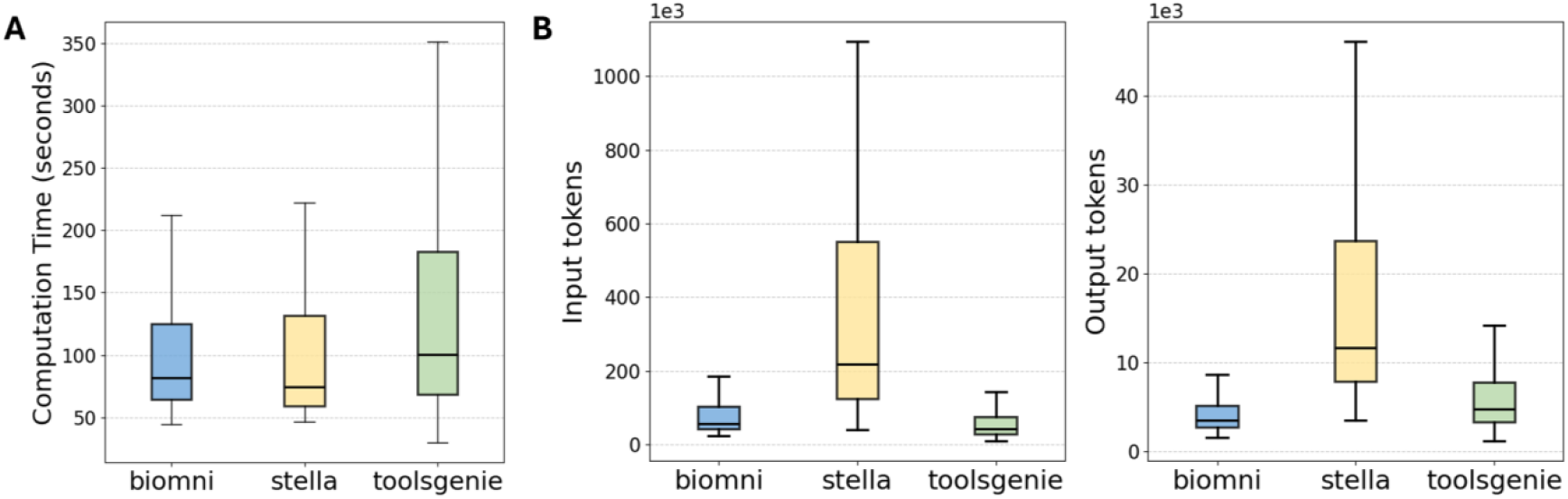
Runtime and token consumption across agents. (A) Box plots of wall-clock computation time per task for each agent. (B) Box plots of input tokens (left) and output tokens (right) consumed per task for each agent.

Token consumption showed more pronounced differences across agents (Figure 4B, Supplementary Table 1b, c). Biomni and ToolsGenie used substantially fewer tokens than STELLA, particularly for input tokens. ToolsGenie generally used fewer input tokens than Biomni but produced moderately more output tokens, consistent with a design that relies less on conditioning over a retrieved biomedical action space and more on generating task-specific executable code during execution. In contrast, STELLA showed markedly higher input and output token usage, with especially long upper tails. This likely reflects the greater context and communication overhead introduced by its manager–subagent coordination, iterative self-refinement, and multi-model orchestration. Together, these patterns highlight the trade-offs each agent makes between execution efficiency and reasoning depth.

## Discussion

The rapid growth of biomedical data has created an urgent need for scalable analytical frameworks that extend beyond traditional approaches dependent on specialized human expertise. LLM-based agents, which can interpret natural-language instructions, generate and execute code, and coordinate multi-step workflows, offer a promising route toward more autonomous, adaptive, and accessible bioinformatics analysis. However, realizing its full potential requires evaluation efforts that assess both successful workflow completion and concordance with expert-generated references in realistic analytical settings. Existing benchmarks have largely focused on question answering, multiple-choice reasoning, or narrowly scoped tasks, and therefore do not fully capture the file-centric, multi-step workflows that characterize routine bioinformatics practice. To address this gap, we developed PromptBio-Bench, a benchmark suite comprising 244 expert-curated tasks across three difficulty levels and spanning diverse areas of bioinformatics and data science. Coupled with a hybrid evaluation framework that integrates format-specific file comparison, quantitative similarity metrics, and LLM-as-judge assessment, PromptBio-Bench enables rigorous end-to-end evaluation of agent performance across heterogeneous analytical outputs.

We evaluated three state-of-the-art bioinformatics agents, Biomni, STELLA, and ToolsGenie, on PromptBio-Bench to assess their ability to execute realistic computational biology and data science tasks. Overall, Biomni and ToolsGenie emerged as comparably strong performers, achieving near-ceiling task completion rates of 0.99 and 0.98, respectively, and the same overall accuracy score of 0.76, whereas STELLA achieved a lower completion rate of 0.88 and a slightly lower accuracy score of 0.72. These results suggest that current agents can often produce valid and evaluable outputs, but that faithfully reproducing expert-generated analyses remains a substantial challenge. This gap became more pronounced as task difficulty increased: on high-difficulty workflows, ToolsGenie and Biomni retained moderate accuracy scores of 0.69 and 0.67, respectively, whereas STELLA declined to 0.56. The difficulty-dependent decline across all agents suggests that complex multi-step workflows requiring coordinated planning, execution, interpretation, and error recovery remain challenging for current agentic systems. The performance profile of STELLA may reflect additional variability associated with its flexible self-evolving architecture, while its higher runtime and token usage are consistent with the overhead introduced by iterative refinement and multi-model orchestration. More broadly, the comparable performance of Biomni and ToolsGenie suggests that robust execution environments, effective tool use, and well-designed workflow scaffolding may be as important as model capability for reliable bioinformatics automation. Future development should therefore focus not only on improving reasoning and domain knowledge, but also on strengthening execution stability, verification mechanisms, cost efficiency, and robustness on high-complexity workflows.

Several limitations of the current study should be acknowledged. Although PromptBio-Bench includes 244 tasks spanning bioinformatics and data science, it remains modest relative to the breadth of modern computational biology, and some task types, data modalities, or analytical scenarios may be underrepresented. Performance estimates within individual subcategories are also based on limited task numbers, which constrains statistical power and cautions against overinterpreting fine-grained differences. In addition, expert reference answers capture one valid analytical path rather than the full range of scientifically acceptable solutions. The evaluation framework may therefore penalize methodologically distinct but valid approaches, particularly for tasks that can be addressed through multiple analytical strategies. Future versions could mitigate this limitation by incorporating multiple reference answers, more flexible equivalence rubrics, and expanded human review for ambiguous cases. The tested configurations may also not fully reflect the potential of all agentic systems. For example, STELLA’s self-evolving feature was not fully utilized, and its results should therefore be interpreted as a lower bound on its capability. We also did not include native LLM coding harnesses, such as Codex and Claude Code; comparing these systems with domain-oriented agentic frameworks would help clarify the relative strengths of bioinformatics-specific agents over general-purpose coding environments. Finally, our single-pass, one-shot protocol does not capture run-to-run variability arising from stochastic LLM generation or the capabilities that may emerge through iterative user feedback. As agentic systems become increasingly integrated into bioinformatics research, future benchmarks should incorporate replicate evaluations and interactive settings that better reflect practical analytical workflows, including iterative refinement of goals, inspection of intermediate outputs, and multi-round analyst–agent interactions.

Despite the limitations, PromptBio-Bench provides a foundation for systematic evaluation of agentic systems in computational biology. Its key resources include a difficulty-stratified, expert-validated task suite spanning diverse bioinformatics and data science domains; a modular evaluation framework that integrates format-specific comparison handlers with LLM-as-judge assessment for heterogeneous outputs; and an extensible, openly available infrastructure designed to support community adoption and continued development. As foundation models and agentic systems continue to evolve and become increasingly integrated into scientific research, rigorous benchmarking of analytical correctness, robustness, and computational efficiency will be essential. PromptBio-Bench is intended to serve as a living resource for the field, providing infrastructure to track meaningful progress and guide the development of AI systems that can reliably support biological discovery.

## Methods

### Benchmark Task Design and Curation

PromptBio-Bench comprises 244 tasks curated to reflect the breadth and analytical complexity of real-world bioinformatics analysis. These tasks were designed and validated by experienced bioinformaticians and data scientists with expertise spanning multi-omics analysis, data science, and machine learning. Each task consists of a natural-language prompt, input files in domain-appropriate formats (e.g., FASTQ, BAM, VCF, CSV), reference answer files prepared by human experts using established tools and pipelines, and an evaluation guideline. Task descriptions were written to reflect the level of specificity typical of requests submitted to a bioinformatics platform, which is specific enough to define the analytical objective but not so prescriptive as to dictate tool choice or parameter settings, unless tool specification was part of the task requirement.

Tasks are organized into two primary domains: Bioinformatics and Data Science, spanning several subfields including sequence data processing, genomics, epigenomics, transcriptomics, proteomics, single-cell and spatial omics, multi-omics integration, data wrangling and visualization, statistical inference, machine learning, survival analysis, among others. Task difficulty was assigned across three tiers using an LLM-ensemble voting procedure, in which each task was independently rated by three AI chat tools: ChatGPT 5.2, Grok 4.1, and Claude 4.6, with ratings repeated three times per tool, and the final label determined by majority vote (Supplementary Figure 3).

### Agent Configuration and Running

We evaluated three general-purpose bioinformatics agents: Biomni (v0.0.8), STELLA (v1.0.0), and ToolsGenie (v3.2.0). All agents were configured with the recommended settings per their official documentation. Biomni and ToolsGenie were configured with Claude Sonnet 4.6 as their LLM backend. STELLA employed its hybrid model strategy as suggested by their paper: the Dev Agent and Tool Creation Agent utilized Claude Sonnet 4.6, while the Manager Agent and Critic Agent were powered by Gemini 2.5 Pro. Each agent was deployed in a Docker image and executed with identical task descriptions and input files. No task-specific prompting, few-shot examples, or agent-specific customization was applied beyond each agent’s default configuration. All agents ran on the same server (Ubuntu 20.04 LTS), with sufficient CPU and RAM allocated across tasks, and a 1-hour runtime limit per task. Runs that exceeded the time limit were killed and recorded as failed.

### Output Evaluation Framework

A central challenge in benchmarking bioinformatics agents is that their outputs span a far wider range of data types than those encountered in question-answering or machine learning benchmarks. Agent outputs may include raw or processed sequence files such as alignment records, variant calls, genomic interval annotations, gene expression matrices, protein structures, or qualitative outputs such as figures and narrative summaries. No single evaluation method is appropriate across this diversity: a metric used in comparing the overlap of genetic variants is meaningless when applied to narrative summary text. To address this problem, we developed a modular evaluation framework comprising format-specific comparison handlers covering major bioinformatics file types. All handlers share a common two-phase architecture: a validation phase that verifies file existence, format compliance via magic-byte signature detection, and file parsability; followed by a comparison phase in which the LLM recommends the most appropriate comparison strategy and parameter configuration based on the file format, question context and evaluation guideline. Only files passing both phases proceed to similarity scoring, and all handlers report results as a normalized similarity score in [0, 1], enabling consistent aggregation across heterogeneous output types.

Each format handler exposes multiple comparison strategies reflecting different tolerance levels and semantic perspectives on similarity. Three tiers are shared across most handlers: *exact* comparison requires record-level identity and is applied when output reproducibility is the explicit evaluation criterion; *approximate* comparison performs record-level matching within a configurable tolerance threshold that accommodates minor, analytically irrelevant variation; and *summary* comparison assesses statistical summaries or distributional agreement between output files without requiring record-by-record details. Several handlers additionally expose specialized strategies where data semantics warrant them, such as a *functional* strategy that validates index files (e.g., .bai, .tbi) for integrity without content comparison, and a *semantic* strategy that delegates image and free-text outputs to a multimodal LLM judge. Full per-format comparison logic and configurable parameters will be described in the Supplementary Note.

### Similarity Metrics

For structured format output, quantitative similarity is computed and normalized using a shared library of metrics, each tailored to the data characteristics and comparison purpose. For set-based comparisons, Jaccard similarity measures the overlap of variants, genomic regions, or clustering labels. For distributional comparisons, Hellinger distance captures agreement between probability distributions such as k-mer frequencies, length distributions, and signal value histograms. For numeric properties such as feature counts, component proportions, and total nucleotide length, Relative Absolute Error (RAE) similarity is applied. For paired numeric vectors such as signal tracks and score columns, Pearson and Spearman correlation assess linear and rank-based agreement, respectively. For sequence-level comparisons, Smith-Waterman normalized alignment identity is applied to nucleotide and protein sequence pairs; for three-dimensional structural comparison, RMSD combined with chain sequence identity is used for protein structures. All metrics map to [0, 1] with 1.0 indicating perfect agreement.

### LLM-as-Judge Evaluation

For unstructured output formats such as figures and free-text summaries, similarity was assessed using an LLM judge (GPT-5.4). For each task, the judge receives the task description, the reference answer file generated by a human expert, and the candidate file produced by the agent. For images, the judge is instructed to assess scientific content equivalence, focusing on whether the same data presentation, patterns, and conclusions are conveyed, rather than pixel-level similarity. For free-text outputs, the judge is instructed to assess whether the key information from the reference is present in the candidate, with particular attention to numerical values. In all cases, the judge returns a structured result comprising an equivalence score on [0, 1], a confidence score, a categorical recommendation (equivalent/partially equivalent/not equivalent/uncertain), and a textual explanation of the recommendation reasoning.

### Task-Level Scoring and Aggregation

Each task yields a single scalar similarity score in [0, 1]. For tasks whose outputs comprise multiple files, the task-level score is the unweighted mean of the individual file similarity scores. Tasks that failed validation, due to failed running, missing outputs, or unparseable files, received a similarity score of 0 and were flagged as non-completions. A task was deemed accurate if its similarity score exceeded the threshold of 0.5, indicating acceptable analytical equivalence with the reference solution. Aggregated accuracy for each agent was then computed within each difficulty stratum, enabling systematic comparison across agents and task categories.

### Computational Cost Assessment

Two dimensions of computational cost were recorded for each agent run: wall-clock execution time and token consumption. Execution time was measured from task submission to the final output. Token consumption was extracted from the API usage logs of each agent’s LLM backend, with input and output tokens recorded separately. For agents using a multi-agent framework, the total token count per task was calculated by summing the input and output tokens across all LLM calls within a single run, respectively.

## Code and Data Availability

The PromptBio-Bench code and data are publicly available. The benchmark evaluation code is available on GitHub at: https://github.com/PromptBio/promptbio-bench. The PromptBio-Bench task capsules, including task descriptions, input data, and reference answers, are available through Hugging Face at: https://huggingface.co/datasets/promptbio-ai/promptbio-bench-data.

## Acknowledgements

Figure 1A was created using BioRender (https://www.biorender.com/) and is acknowledged accordingly.

## Supplementary Materials

**Supplementary Figure 1.**
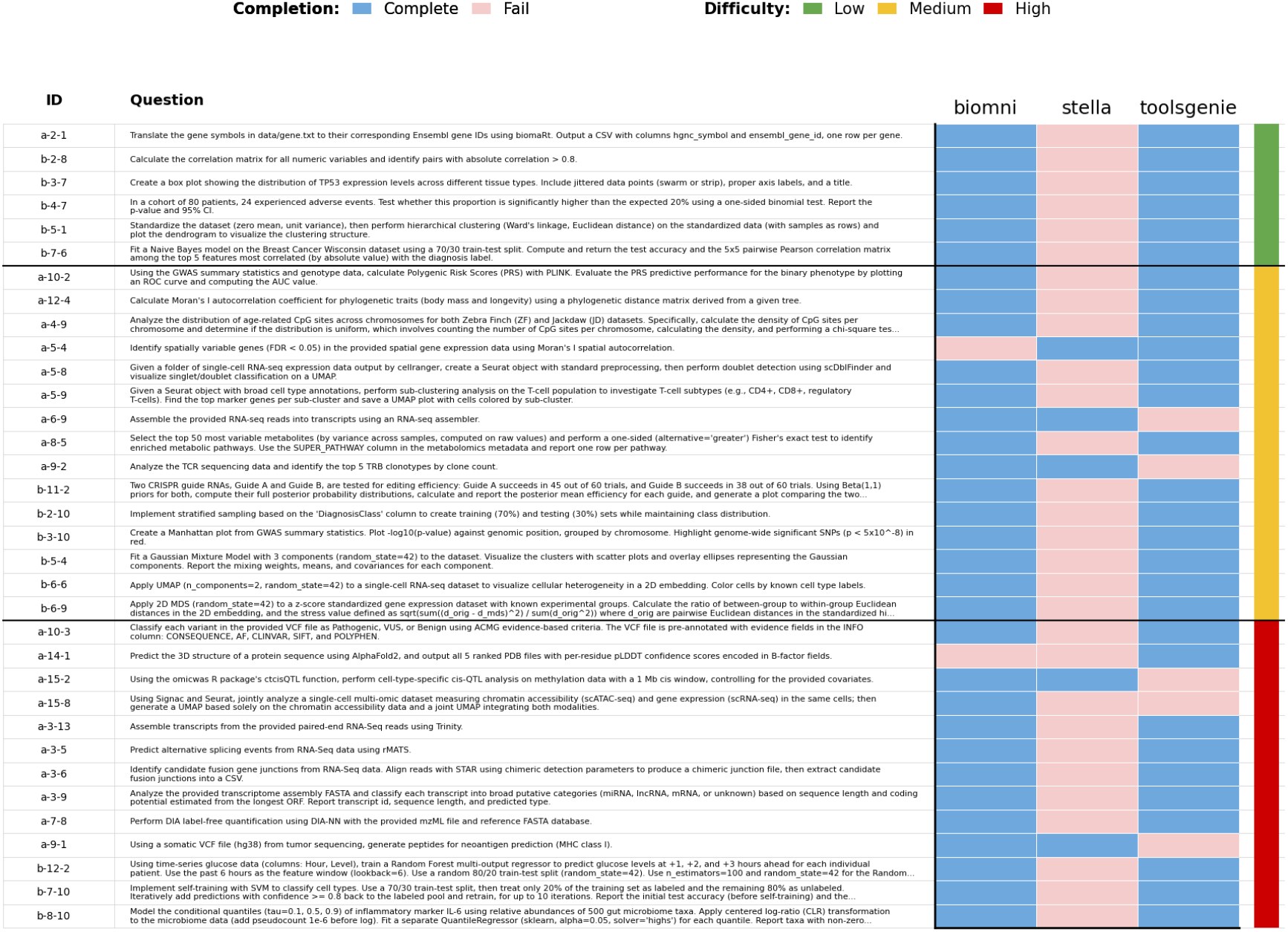
Task failure across agents. Heatmap showing failed tasks across three agents. Blue indicates successful completion, pink indicates failure, and the rightmost color bar denotes task difficulty.

**Supplementary Figure 2.**
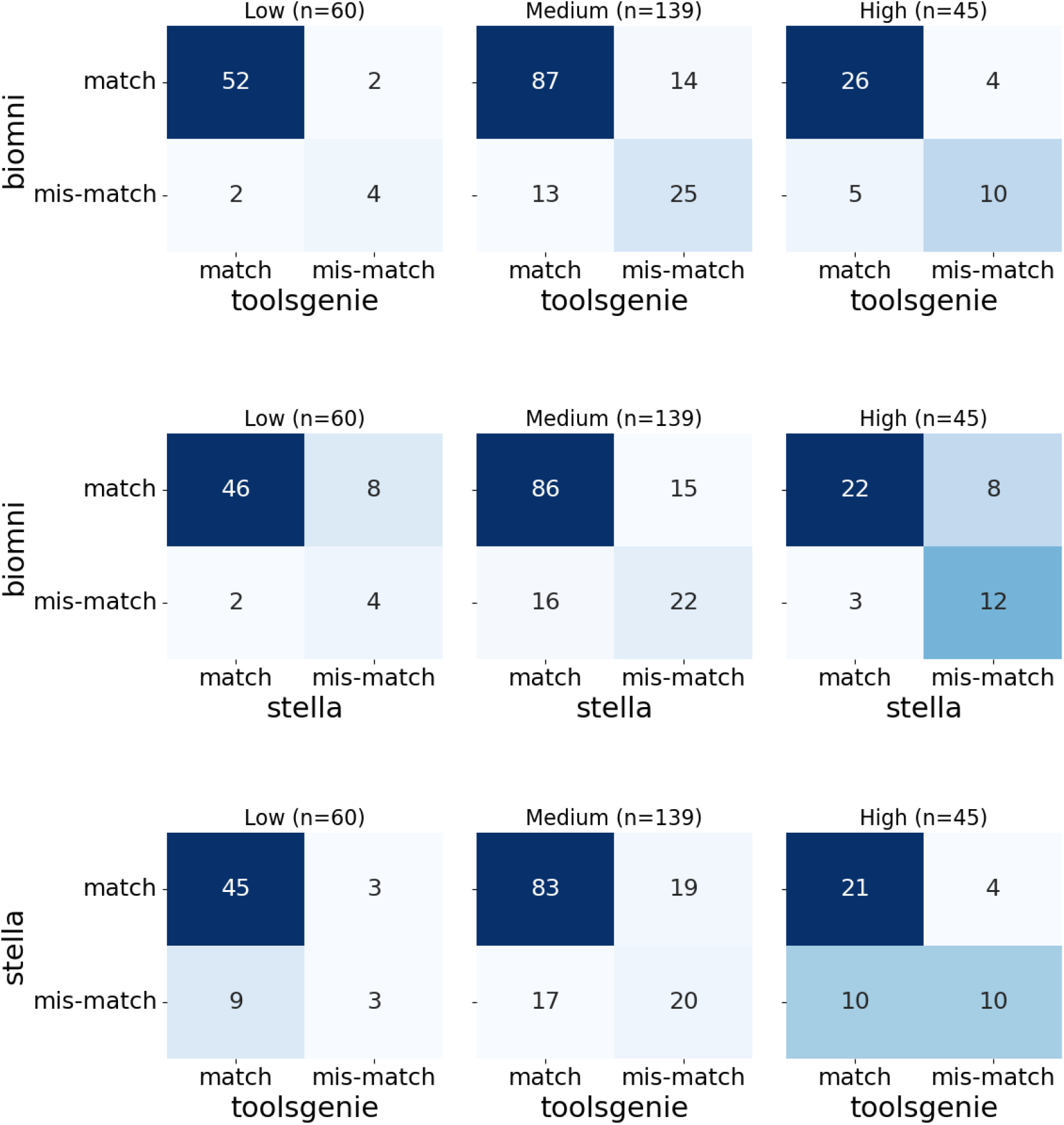
Pairwise confusion matrices comparing agent correctness against the reference answer, stratified by question difficulty. Each 2×2 matrix shows the number of questions in which two agents’ outputs both matched with the reference answer, both mismatched with the reference answer, or differed in their correctness relative to the reference answer, across low, medium, and high-difficulty levels. Rows and columns represent each agent’s outcome (match or mismatch) relative to the reference answer. Darker cells indicate higher counts.

**Supplementary Figure 3.**
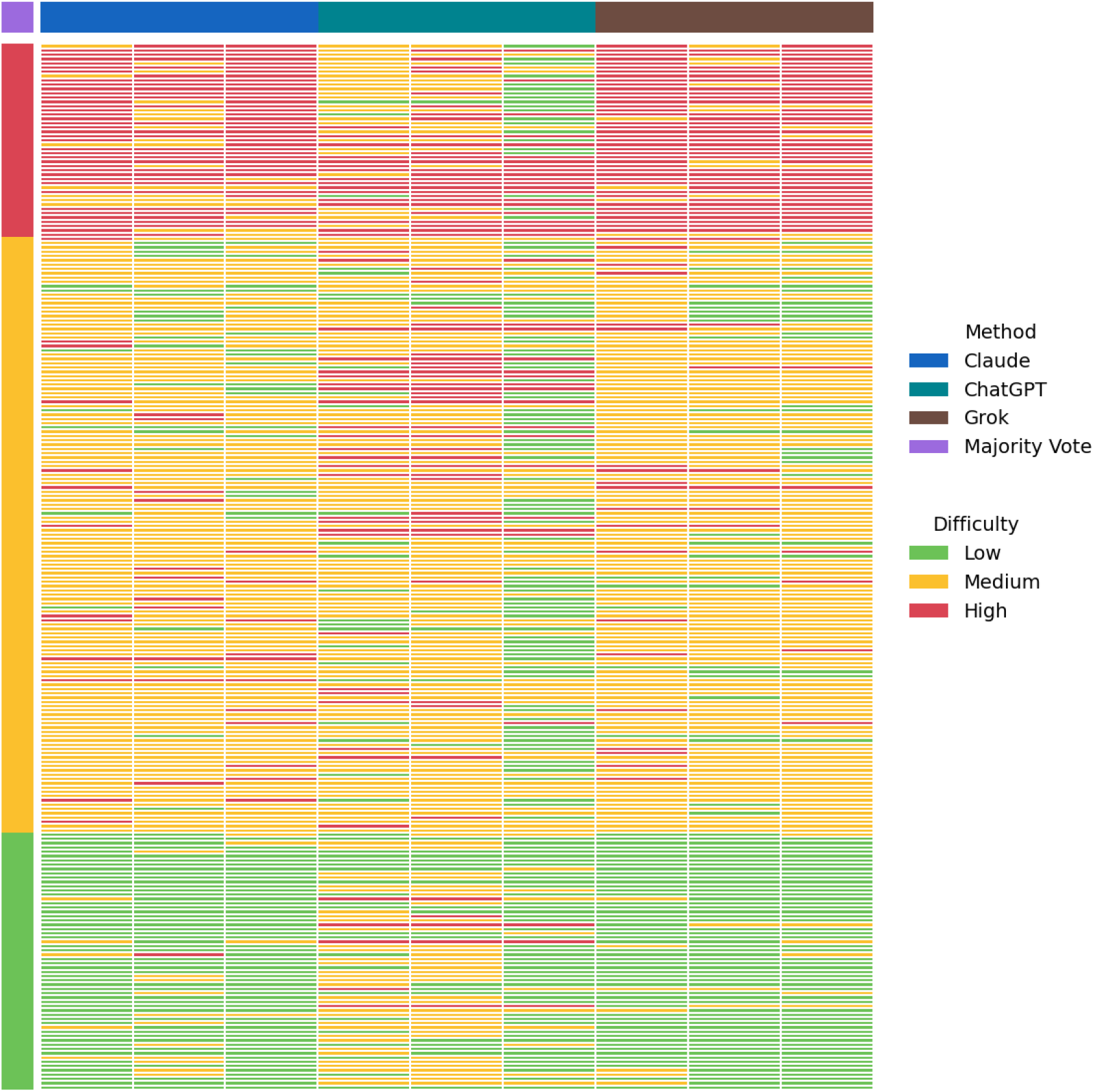
Difficulty level heatmap of benchmark questions. Each column represents an independent classification run (three runs per LLM method), and each row corresponds to a single benchmark question. The final difficulty label was determined by majority vote across all nine runs. Questions are grouped along the y-axis according to their final assigned label.

**Supplementary Table 1:** Computational cost summary for Biomni, STELLA, and ToolsGenie.

(a) Wall-clock (seconds)
| **Agent** | **Min** | **Q25** | **Median** | **Q75** | **Max** | **Mean** | **Std Dev** |
| --- | --- | --- | --- | --- | --- | --- | --- |
| biomni | 44.05 | 64.60 | 81.52 | 124.70 | 3600.18 | 170.51 | 412.34 |
| stella | 46.44 | 59.23 | 74.35 | 131.94 | 3600.21 | 275.25 | 648.58 |
| toolsgenie | 29.62 | 68.41 | 100.48 | 183.05 | 3600.24 | 217.97 | 460.74 |

(b) Input token consumption
| **Agent** | **Min** | **Q25** | **Median** | **Q75** | **Max** | **Mean** | **Std Dev** |
| --- | --- | --- | --- | --- | --- | --- | --- |
| biomni | 23,925 | 41,449 | 54,948 | 102,319 | 934,543 | 98,584 | 122,498 |
| stella | 40,463 | 124,388 | 218,882 | 532,646 | 38,694,347 | 1,294,062 | 3,833,468 |
| toolsgenie | 9,405 | 27,863 | 40,999 | 73,738 | 695,525 | 59,418 | 57,426 |

(c) Output token consumption
| **Agent** | **Min** | **Q25** | **Median** | **Q75** | **Max** | **Mean** | **Std Dev** |
| --- | --- | --- | --- | --- | --- | --- | --- |
| biomni | 1,544 | 2,773 | 3,466 | 5,118 | 41,981 | 4,604 | 3,634 |
| stella | 3,536 | 7,843 | 11,696 | 23,312 | 565,609 | 23,670 | 46,374 |
| toolsgenie | 1,168 | 3,288 | 4,772 | 7,804 | 89,216 | 6,989 | 7,629 |

### Supplementary Note

This note documents each format handler’s supported comparison strategies, configurable parameters, primary similarity metric, and recommended use cases.

#### S1.1 FASTA

FASTA stores named nucleotide or protein sequences (‘>‘-prefixed header followed by sequence lines). Gzip-compressed variants (.fa.gz) are supported. Output equivalence ranges from exact sequence identity (deterministic assembly) to distributional agreement (de novo assemblies produced by different tools).

| Strategy | Description & Parameters | Primary Metric | When to Use |
| --- | --- | --- | --- |
| **exact** | Match by sequence content (default) or by name (*by_name* = True). Parameters: *by_name* (bool, default: False); *ignore_case* (bool, default: True); *single_line* (bool, default: False) — when True, multiline candidate sequences are excluded. | $\text{matched\_seqs} / \text{total\_ref\_seqs} \in [0, 1]$ | Identical sequence output required (e.g., genome assembly reproducibility, format conversion). |
| **approximate** | Greedy content-based pairing; BioPython local (Smith–Waterman) alignment per pair. Parameters: *by_name* (bool, default: False); *ignore_case* (bool, default: True); *single_line* (bool, default: False). | *Mean pairwise alignment identity* $\in [0, 1]$ | Sequences may be nearly identical (assembly polishing, consensus calling). |
| **summary**<br>(default) | Four components: sequence count (RAE), total length (RAE), length distribution (Hellinger), k-mer distribution (Hellinger). Parameters: *k* (int, default: 11); *num_bins* (int, default: 10). | *Mean of 4 component similarities* $\in [0, 1]$ | Large assemblies or metagenomics where record-level comparison is impractical. |

#### S1.2 FASTQ

FASTQ pairs each sequence with per-base Phred quality scores. It is the common output of trimming, filtering, and demultiplexing tools. Minor end-trimming differences between tools motivate the tolerance parameters of the approximate strategy.

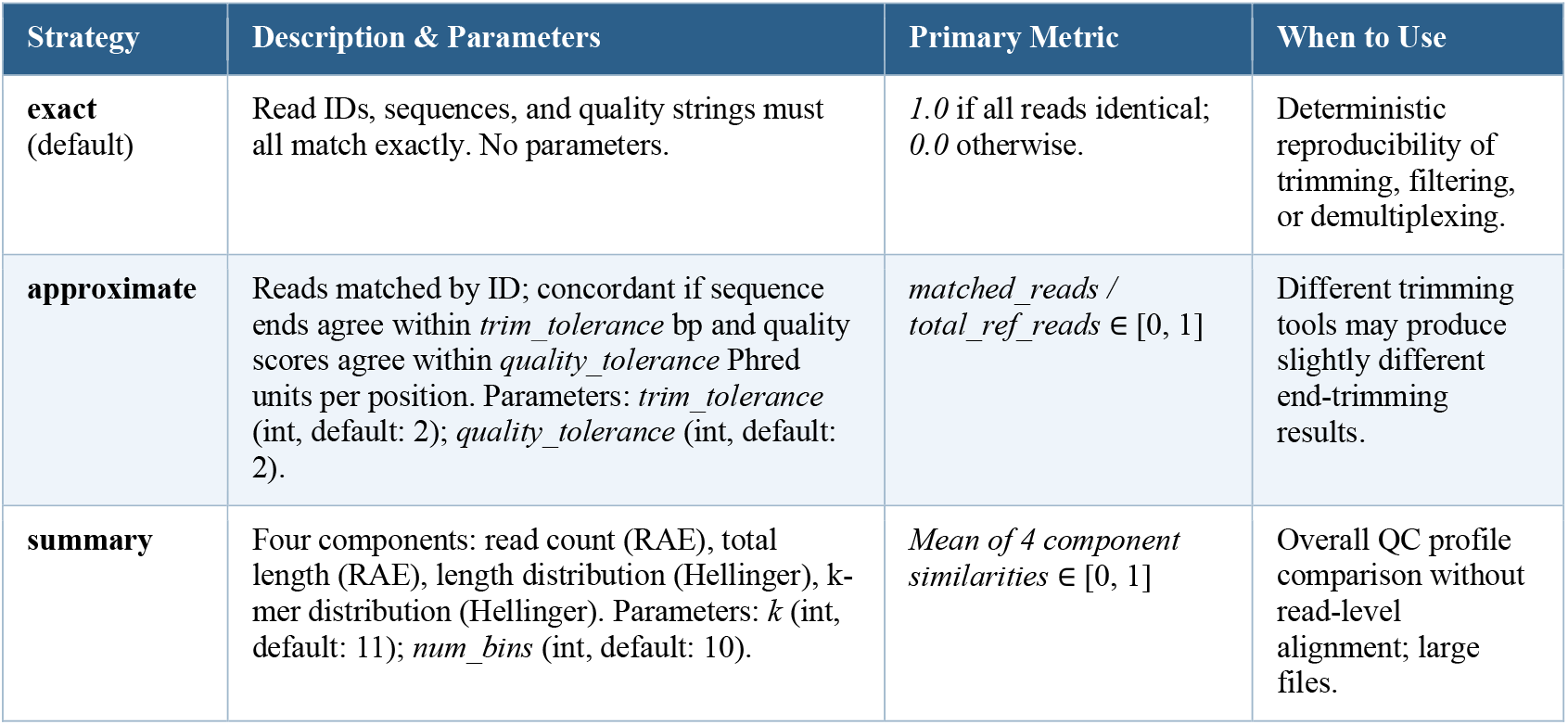

#### S1.3 VCF / BCF

VCF/BCF records variants (SNVs, indels, SVs) with CHROM, POS, REF, ALT, QUAL, FILTER, and INFO/FORMAT fields. Because callers may represent the same variant differently (allele splitting, indel alignment), set-level comparison is preferable for cross-tool evaluation.

| Strategy | Description & Parameters | Primary Metric | When to Use |
| --- | --- | --- | --- |
| **exact** | Ordered variant sets must match exactly (CHROM, POS, REF, ALT, QUAL, FILTER). No parameters. | *1.0* if identical; *0.0* otherwise. | Deterministic variant caller reproducibility. |
| **approximate** | Variants matched as key sets (CHROM, POS, REF, ALT); Jaccard, precision, and recall computed. Parameters: *position_window* (int, default: 10) bp; *require_allele_match* (bool, default: True). | *Jaccard similarity* $\in [0, 1]$; precision and recall reported in details. | Comparing calls from different tools or parameters. |
| **summary**<br>(default) | bcftools stats profiles compared via element-wise RAE across SN, TSTV, AF, and QUAL sections. No parameters. | *Mean RAE similarity over SN section fields* $\in [0, 1]$ | Quick variant-calling quality comparison; default for most VCF tasks. |

#### S1.4 BED / narrowPeak / broadPeak / bedGraph / bigBed

BED stores genomic intervals; variants include narrowPeak/broadPeak (ENCODE peaks), bedGraph (per-interval signal), and bigBed (binary indexed). Five strategies assess output at different granularities: individual regions, base-level coverage, or score patterns.

| Strategy | Description & Parameters | Primary Metric | When to Use |
| --- | --- | --- | --- |
| **exact** | Element-by-element comparison after sorting. Parameters: *coord_only* (bool, default: True); *ignore_order* (bool, default: False); *merge_overlap* (bool, default: False). | *1.0* if all regions match; *0.0* otherwise. | Identical interval output required. |
| **approximate**<br>(default) | Region-level Jaccard: each reference region matched to candidates with $\geq \text{overlap\_threshold}$ overlap. Parameters: *overlap_threshold* (float, default: 0.95); *tolerance* (int, default: 0) bp; *coord_only* (bool, default: True). | *Jaccard similarity* (matched / union regions) $\in [0, 1]$ | Peak callers may produce slightly different boundaries for the same genomic features. |
| **overlap** | Base-level Jaccard: intervals merged, then intersected at bp level via pybedtools. Parameters: *coord_only* (bool, default: True); *merge_overlap* (bool, default: True). | *Base-pair Jaccard* $\in [0, 1]$; precision and recall in details. | Total genomic coverage agreement matters more than individual region boundaries. |
| **correlation** | Pearson or Spearman correlation of score values from overlapping region pairs. Parameters: *overlap_threshold* (float, default: 0.95); *correlation_method* ('pearson' \| 'spearman', default: 'pearson'). | $\vert \text{correlation}\vert \times \text{coverage\_ratio} \in [0, 1]$ | Quantitative score column comparison (ChIP-seq enrichment, signal intensity). |
| **summary** | Aggregate comparison: region count (RAE), average length (RAE), length distribution (Hellinger); optional score distribution (Hellinger). Parameters: *num_bins* (int, default: 50). | *Length distribution Hellinger similarity* $\in [0, 1]$ | Quick aggregate comparison; large interval files. |

#### S1.5 BAM / SAM / CRAM

SAM/BAM/CRAM stores read alignments with 11 mandatory fields (QNAME, FLAG, RNAME, POS, MAPQ, CIGAR, RNEXT, PNEXT, TLEN, SEQ, QUAL) plus optional tags. The approximate strategy uses chunk-based QNAME-sorted streaming to handle large files efficiently.

| Strategy | Description & Parameters | Primary Metric | When to Use |
| --- | --- | --- | --- |
| **exact** | All 11 mandatory fields and common optional tags compared record-by-record. Headers compared excluding @PG and SO fields. Early-exit flagstat check applied first. No parameters. | *1.0* if all records identical; *0.0* otherwise. | Deterministic aligner reproducibility; bit-identical pipeline verification. |
| **approximate** | QNAME-matched read pairs checked: same RNAME/POS, $\text{MAPQ} \geq \text{min\_mapq}$, positional overlap $\geq \text{overlap\_threshold}$. Files auto-sorted by QNAME if needed. Parameters: *min_mapq* (int, default: 20); *overlap_threshold* (float, default: 0.95); *chunk_size* (int, default: 100,000). | *Jaccard similarity* (concordant / union read names) $\in [0, 1]$ | Different aligners or parameters; checks concordance of mapped read placement. |
| **summary**<br>(default) | Flagstat count vectors compared via per-metric RAE; MAPQ distributions via Hellinger distance. No parameters. | *Mean of flagstat RAE and MAPQ Hellinger similarities* $\in [0, 1]$ | Quick alignment quality comparison; default for most BAM tasks. |
| **coverage** | samtools depth -a on both files; per-position depth ratio min/max. Parameters: *depth0_similarity* (float, default: NaN) — score assigned when both depths are zero. | *Mean per-position depth ratio* $\in [0, 1]$ | Coverage uniformity or depth agreement (not read-placement concordance). |
| **variant** | bcftools mpileup \| bcftools call generates a VCF from each BAM; delegates to VCFHandler. Parameters: *ref_fasta* (str, required); *vcf_strategy* (str, default: 'summary'). | *VCF handler score under vcf_strategy* $\in [0, 1]$ | Functional equivalence at the variant level. |

#### S1.6 FASTA Index (.fai)

FASTA index files (.fai) store a five-column table (name, length, byte offset, linebases, linewidth) enabling O(1) sequence access. Byte offsets differ for reformatted FASTAs; the approximate strategy therefore focuses exclusively on the length column.

| Strategy | Description & Parameters | Primary Metric | When to Use |
| --- | --- | --- | --- |
| **exact**<br>(default) | All five fields compared for all sequences (DataFrame equality). No parameters. | *1.0* if DataFrames exactly equal; *0.0* otherwise. | Same FASTA indexed with the same tool; exact reproducibility required. |
| **approximate** | Length column compared via RAE; missing sequences assigned *max_rae*. Parameters: *tolerance* (float, default: 0.0); *max_rae* (float, default: 1e6). | *Proportion of sequences within tolerance* $\in [0, 1]$ | Different indexing tools may produce different offsets or line-widths. |
| **summary** | Three components: sequence count (RAE), total length (RAE), length distribution (Hellinger). Parameters: *num_bins* (int, default: 10). | *Mean of 3 component similarities* $\in [0, 1]$ | Quick check without field-level comparison. |

#### S1.7 BAM / CRAM Index (.bai, .crai)

BAM/CRAM index files (.bai/.crai) encode byte offsets for random-access to genomic regions. The functional strategy compares samtools idxstats output (per-reference read counts) rather than binary content, which may vary between index versions.

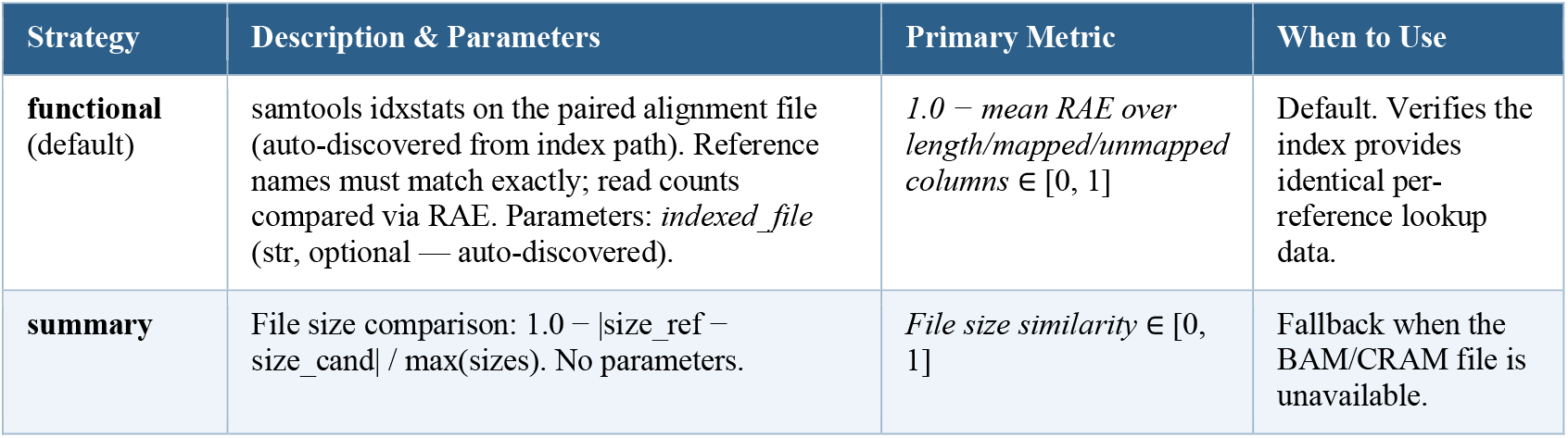

#### S1.8 Tabix Index (.tbi)

Tabix index files (.tbi) enable random access into bgzip-compressed VCF, BED, and GTF files. The functional strategy compares indexed contig names and lengths rather than binary content.

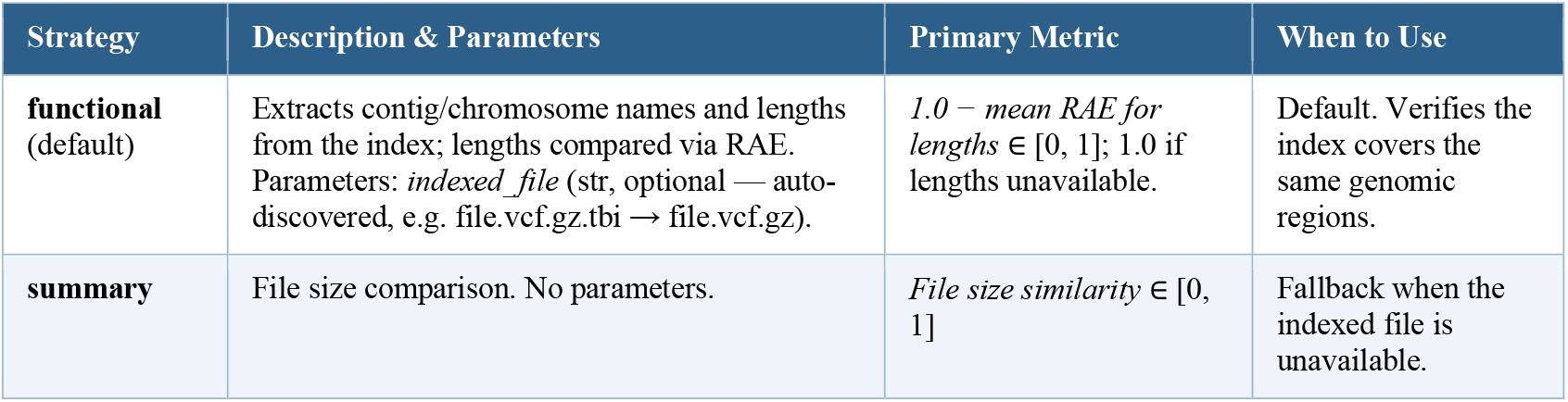

#### S1.9 bigWig / WIG

bigWig/WIG stores continuous genomic signal tracks (read depth, ChIP-seq enrichment, methylation) as per-position floating-point values. Tracks may differ in absolute scale while preserving the same pattern, motivating the scale-invariant correlation strategy.

| Strategy | Description & Parameters | Primary Metric | When to Use |
| --- | --- | --- | --- |
| **exact** | Chromosome names/sizes must match exactly; all position values must be identical. Parameters: *max_positions* (int, default: None). | 1.0 if all values match; 0.0 otherwise. | Identical signal track output required. |
| **approximate** | Fraction of positions within relative tolerance; weighted by chromosome size × coverage ratio. Parameters: *tolerance* (float, default: 1.0); *max_positions* (int, default: None). | *weighted_similarity* × *coverage_ratio* $\in [0, 1]$ | Signal values may differ slightly due to normalisation or rounding. |
| **correlation** | Pearson or Spearman correlation per chromosome; weighted by size × coverage ratio. Scale-invariant. Parameters: *correlation_method* ('pearson' \| 'spearman', default: 'pearson'); *max_positions* (int, default: None). | *weighted_\|correlation\|* × *coverage_ratio* $\in [0, 1]$ | Same signal pattern with different absolute scale (e.g., different normalisation methods). |
| **summary**<br>(default) | Per-chromosome value histograms compared via Hellinger distance; weighted by size × coverage ratio. Parameters: *max_positions* (int, default: None); *num_bins* (int, default: 50). | *weighted_Hellinger_similarity* × *coverage_ratio* $\in [0, 1]$ | Quick signal distribution comparison; default for most bigWig tasks. |

#### S1.10 PDB / mmCIF

PDB/mmCIF stores three-dimensional atomic coordinates in chains and residues. These formats are produced by structure determination (X-ray crystallography, cryo-EM) and prediction tools (AlphaFold2, ESMFold). Chains may differ in identifier or completeness; sequence alignment therefore precedes structural comparison.

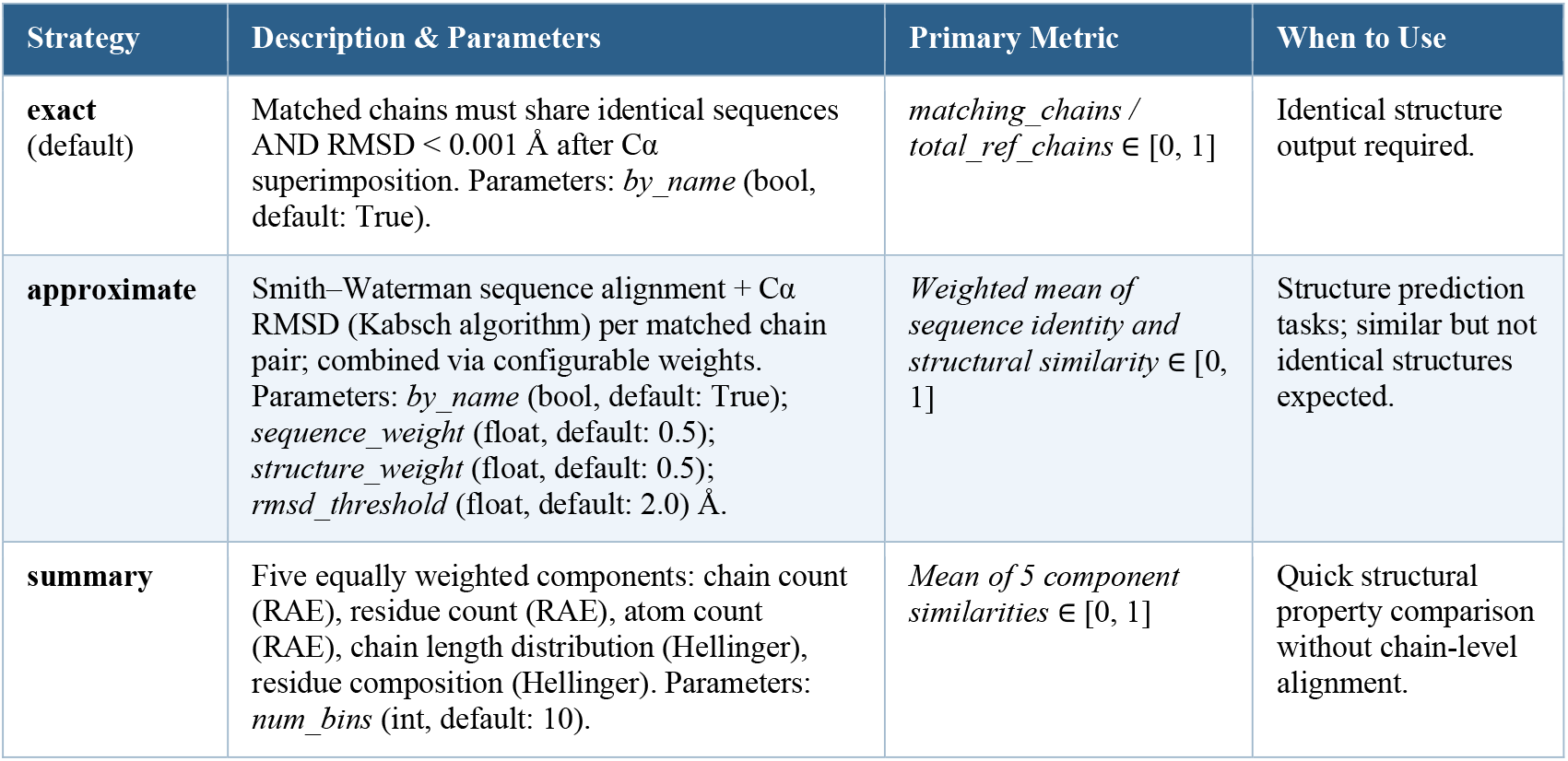

| Strategy | Description & Parameters | Primary Metric | When to Use |
| --- | --- | --- | --- |
| **exact**<br>(default) | Matched chains must share identical sequences AND RMSD $< 0.001$ Å after Ca superimposition. Parameters: *by_name* (bool, default: True). | *matching_chains* / *total_ref_chains* $\in [0, 1]$ | Identical structure output required. |
| **approximate** | Smith–Waterman sequence alignment + Ca RMSD (Kabsch algorithm) per matched chain pair; combined via configurable weights. Parameters: *by_name* (bool, default: True); *sequence_weight* (float, default: 0.5); *structure_weight* (float, default: 0.5); *rmsd_threshold* (float, default: 2.0) Å. | *Weighted mean of sequence identity and structural similarity* $\in [0, 1]$ | Structure prediction tasks; similar but not identical structures expected. |
| **summary** | Five equally weighted components: chain count (RAE), residue count (RAE), atom count (RAE), chain length distribution (Hellinger), residue composition (Hellinger). Parameters: *num_bins* (int, default: 10). | *Mean of 5 component similarities* $\in [0, 1]$ | Quick structural property comparison without chain-level alignment. |

#### S1.11 CSV / TSV / XLSX

CSV/TSV/XLSX are standard output formats for differential expression, enrichment, clustering, and summary statistics. Equivalent tables may differ in column names, row order, or numeric precision. The approximate strategy employs LLM-assisted alignment to handle these structural differences before cell-level comparison.

| Strategy | Description & Parameters | Primary Metric | When to Use |
| --- | --- | --- | --- |
| **exact** | Cell-by-cell equality (`DataFrame.equals()`); numeric tolerance configurable. Parameters: *float_tolerance* (float, default: 1e-9). | *matching_cells* / *total_cells* $\in [0, 1]$ | Identical tabular output required; rarely appropriate for bioinformatics. |
| **approximate**<br>(default) | Three-phase pipeline: (1) LLM plans alignment (transpose, column mapping, row IDs); (2) deterministic structural alignment; (3) LLM-planned per-column comparison (Pearson/Spearman/RAE for numeric; exact/overlap for categorical). Parameters: *question* (str, default: 'Compare two tabular data files for equivalence'). | *Mean per-column similarity scores* $\in [0, 1]$ | Tables may differ in column names, row order, or numeric precision. Recommended default. |
| **summary** | No LLM calls. Shape, column Jaccard, missing-value rate, numeric distribution (Hellinger), | *Mean of structural/distributional* | Large tables where LLM calls are too |
| | categorical distribution (Hellinger). Mean of components. No parameters. | *component similarities* $\in [0, 1]$ | costly; only aggregate statistics matter. |
| **semantic** | Both tables (up to *max_rows* rows) submitted to a multimodal LLM for holistic equivalence judgment. Parameters: *question* (str); *max_rows* (int, default: 1,000). | *LLM equivalence score* $\in [0, 1]$ | Different representations of the same biological finding; semantic understanding required. |

#### S1.12 Image

Supported formats: PNG, JPEG, TIFF, BMP, GIF, WEBP, SVG, and PDF. SVG and PDF files are rasterised prior to submission. Because two correct figures may differ in colour or layout while conveying identical biological content, pixel-level comparison is generally uninformative.

| Strategy | Description & Parameters | Primary Metric | When to Use |
| --- | --- | --- | --- |
| **semantic** (default) | SVG→PNG via cairosvg; PDF→PNG via pdf2image. Images base64-encoded and submitted to a multimodal LLM with task context. Assesses scientific content equivalence, not pixel similarity. Parameters: *question* (str, optional); *model* (str, default: 'gpt-5.4'). | *LLM equivalence score* $\in [0, 1]$; confidence and explanation reported in details. | All image outputs. Only available strategy. |

#### S1.13 Plain Text

Plain text files (.txt) contain analysis summaries, QC reports, log files, or numeric results. Strategy selection depends on the expected type of variation: structured logs are best handled by the approximate strategy, narrative summaries by the semantic strategy, and files containing a single scalar value by the numeric strategy.

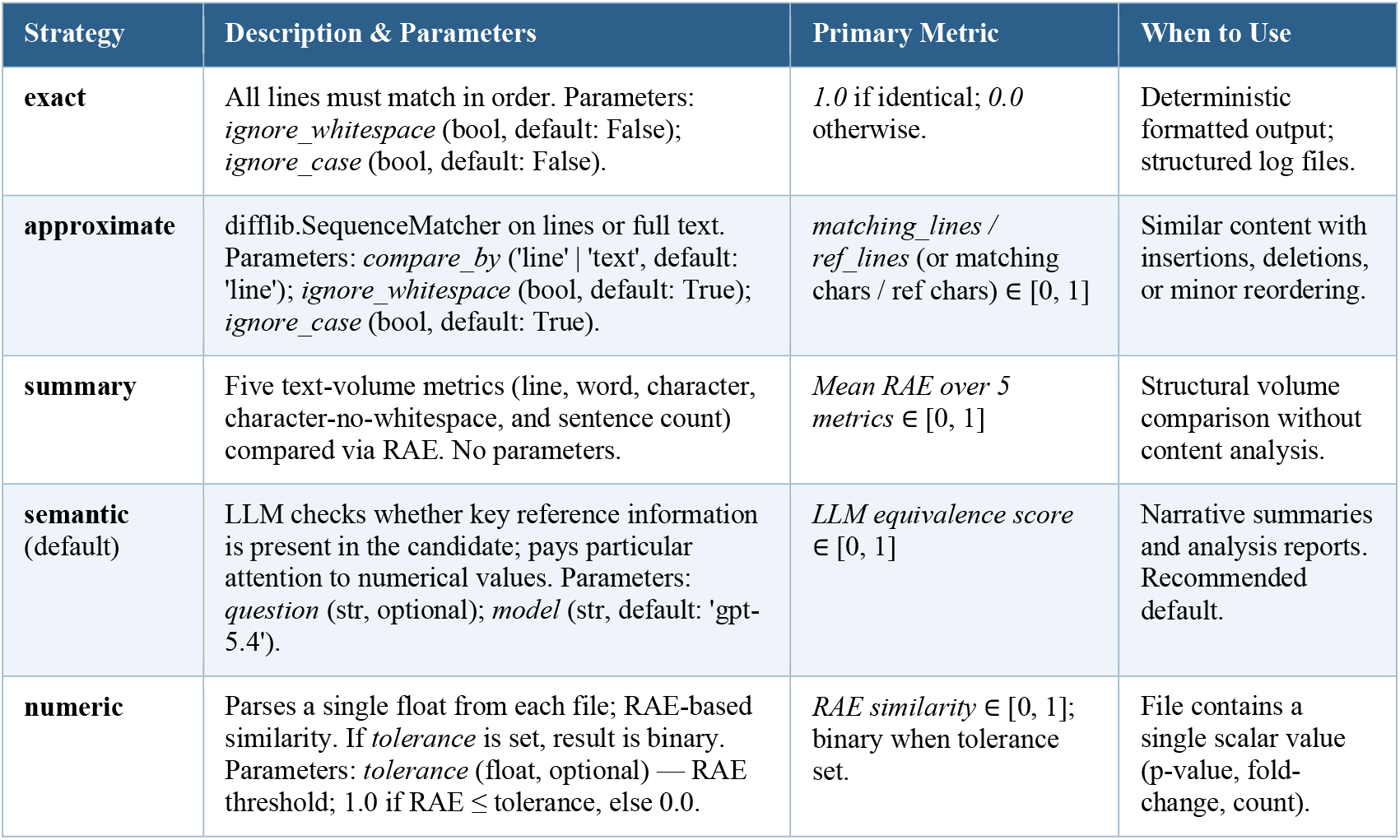

